# Leveraging the resting brain to predict memory decline after temporal lobectomy

**DOI:** 10.1101/2022.12.22.521602

**Authors:** Sam Audrain, Alexander Barnett, Mary Pat McAndrews

## Abstract

**Objectives:** Anterior temporal lobectomy as a treatment for temporal lobe epilepsy is associated with a variable degree of postoperative memory decline, and estimating this decline for individual patients is a critical step of preoperative planning. Presently, predicting memory morbidity relies on indices of preoperative temporal lobe structural and functional integrity. However, epilepsy is increasingly understood as a network disorder, and memory a network phenomenon. We aimed to assess the utility of functional network measures to predict postoperative memory changes.

**Methods:** Patients with left and right temporal lobe epilepsy (TLE) were recruited from an epilepsy clinic. Patients underwent preoperative resting-state fMRI (rs-fMRI) and pre- and postoperative neuropsychological assessment approximately one year after surgery. We compared functional connectivity throughout the memory network of each patient to a healthy control template based on 19 individuals to identify differences in global organization. A second metric indicated the degree of integration of the to-be-resected temporal lobe with the rest of the memory network. We included these measures in a linear regression model alongside standard clinical and demographic variables as predictors of memory change after surgery.

**Results:** Seventy-two adults with TLE were included in this study (37 left/35 right). Left TLE patients with more abnormal memory networks, and with greater functional integration of the to-be-resected region with the rest of the memory network preoperatively, experienced the greatest decline in verbal memory after surgery. Together, these two measures explained 44% of variance in verbal memory change (F(2,31)=12.01, p=0.0001), outperforming standard clinical and demographic variables. None of the variables examined in this study were associated with visuospatial memory change in patients with right TLE.

**Conclusion:** Resting-state connectivity provides valuable information concerning both the integrity of to-be-resected tissue as well as functional reserve across memory-relevant regions outside of the to-be-resected tissue. Intrinsic functional connectivity has the potential to be useful for clinical decision-making regarding memory outcomes in left TLE, and more work is needed to identify the factors responsible for differences seen in right TLE.

## Introduction

While anterior temporal lobe (ATL) resections are quite effective at seizure management in patients with temporal lobe epilepsy (TLE)^1^, left anterior temporal lobectomies confer the risk of a variable degree of verbal memory and naming decline, and resections in the right hemisphere are often associated with postoperative visuospatial memory deficits^1,2^. Developing techniques to accurately predict postoperative cognitive morbidity after surgery is critical to the clinical care of TLE patients. Presently, methods of predicting cognitive decline for a given patient rely largely on indices of preoperative functional adequacy of temporal lobe tissue. Patients who perform well on preoperative neuropsychological tests of material-specific memory (verbal memory in patients with left TLE and visuospatial memory in patients with right TLE)^2,3^, who have structurally intact medial temporal lobes^4,5^, and who show greater functional activation in the to-be-resected medial temporal lobe during mnemonic processing, as measured with task-based fMRI, all tend to exhibit the greatest decline in memory after the surgery^6–8^. Together these methods agree, perhaps predictably, that resecting temporal lobe tissue that is functionally healthy is detrimental to memory.

While it is clear that focal indices of temporal lobe functioning are clinically useful, TLE is increasingly considered a network disorder^9,10^, and memory a network phenomenon^11^. Some degree of functional network reorganization has been described in patients with TLE^12–14^, and several studies using resting-state fMRI (rs-fMRI) have reported an association between preoperative memory capacity and connectivity between memory-relevant regions in this population^12,15–21^. Coupling within memory networks at rest in TLE therefore appears to be functionally significant^10,18^, and could capture functional reserve untapped by metrics of temporal lobe integrity.

It follows that there is potential for resting-state network measures to add predictive value to the clinical decision-making process regarding memory outcomes. Indeed, the fact that resting-state scans can be collected in mere minutes, place no cognitive demands on the patient, and can be used to examine multiple cognitive networks with a single acquisition, positions rs-fMRI as an attractive tool for use in the clinic^22^. With a single scan, it is possible to evaluate multiple metrics of brain organization that range from relatively focal indices of connectivity of the epileptogenic region to broader network organization across distributed brain regions, which may each explain unique variance in cognitive change after surgery. Finally, it is unclear whether altered coupling between memory regions supports or hinders functional resilience, and so consideration of broader network connectivity and topology is needed to understand the brain’s capacity to withstand network insult from surgery.

Here, we sought to evaluate the utility of preoperative resting-state network connectivity to predict postoperative material-specific memory change in patients with left and right TLE. To do so, we evaluated two rs-fMRI metrics of the memory network: one that broadly captures memory network organization relative to the ‘normative’ pattern of controls (termed ‘matrix similarity’), and a more focal measure (‘mean ATL degree’) that captures functional integration between to-be-resected tissue and the rest of the memory network. We previously found that these complimentary metrics, as applied to the preoperative resting-state language network, explained unique variance in naming change after ATL resection in patients with left TLE^23^. In line with that work, we hypothesized that preoperative memory network deviation from the normative pattern and greater integration of the to-be-resected ATL with the rest of the memory network would associate with decline in material-specific memory after surgery. To contextualize the utility of these metrics in the prediction of memory change, we additionally compared them to standard clinical and demographic variables associated with postoperative memory outcomes in the literature (age, sex, education, age of onset, presence of medial temporal sclerosis (MTS), and preoperative memory scores)^24–26^.

## Materials and Methods

### Participants

Seventy-two English-speaking adults with medically intractable TLE participated in this study. Thirty-seven patients were diagnosed with left TLE and 35 with right TLE according to continuous video-EEG recordings (see **Table 1** for demographic details). Sixty-four patients (34 LTLE, 30 RTLE) subsequently underwent a standard ATL resection, which involved the removal of approximately 1cm from the superior temporal gyrus, 3cm from the middle temporal gyrus, and 4-5cm from the inferior temporal gyrus extending caudal from the temporal pole^27^. The entire hippocampus and amygdala were also removed. Eight patients (3 LTLE, 5 RTLE), underwent a selective amygdalohippocampectomy rather than a standard ATL resection where the entire hippocampus and amygdala were removed, leaving the temporal neocortex largely intact^27^. All patients who underwent surgery achieved good postoperative seizure outcomes at one year (Engel scores 1a-d). Nineteen healthy control subjects with no history of neurological or psychiatric disorder were also recruited for rs-fMRI. All data were collected between the years of 2008-2018 at the Epilepsy Clinic at Toronto Western Hospital. Sample size was determined based on the number of eligible complete cases in that 10-year time-period. Our previous work using the same methodology indicates our sample size, which is larger in the present manuscript, is sufficient for this type of analysis.^23^

**Table 1.** Demographic data.

|  | **Controls** | **Left TLE** | **Right TLE** |
| --- | --- | --- | --- |
| *N* | 19 | 37 | 35 |
| Preoperative Age, y (SD) | 34.42 (10.72) | 35.84 (11.66) | 38.43 (10.95) |
| Education, y (SD) | 18.01 (3.10) | 14.55 (2.25) | 14.37 (2.72) |
| Sex, M/F | 11/8 | 14/23 | 13/22 |
| Preoperative material-specific memory, *z* (SD) | – | -0.44 (1.4) | -0.10 (0.99) |
| Postoperative material-specific memory change, *z* (SD) | – | -0.33 (1.11) | -0.04 (0.92) |
| Language Dominance, L/R/B/NA | – | 31/0/6/0 | 30/0/3/2 |
| Age of Onset, y (SD) | – | 16.89 (14.17) | 20.01 (13.30) |
| Presence of MTS, yes/no | – | 27/10 | 23/12 |
| Surgery Type, ATL/sAH | – | 34/3 | 30/5 |

ATL = standard anterior temporal lobectomy; B = bilateral; F = female; L = left; M = male; MTS = mesial temporal sclerosis; NA = not available; R = right; sAH = selective amygdalohippocampectomy; SD = standard deviation; TLE = temporal lobe epilepsy; y = years, *z* = *z*-score. Language dominance was determined by task-based fMRI using standard activation paradigms involving covert word generation. Characterization of MTS was based on radiology (3T MRI protocol).

### Standard Protocol Approvals, Registrations, and Consents

This study was approved by the University Health Network Ethics Board. Patients either gave prospective written informed consent, or, for an older cohort, permission for retrospective analysis of clinical data was obtained from the University Health Network Ethics Board. All controls gave prospective written informed consent.

### Neuropsychological testing

A comprehensive battery of neuropsychological tests was administered to patients preoperatively and at a postoperative session approximately 12 months after surgery. In the present study, we focused on verbal memory in patients with left TLE, and visuospatial memory in patients with right TLE, given the typical pattern of material-specific memory loss in these cohorts^3^. Summary verbal and visuospatial memory scores were calculated based on a previously reported principal component analysis from our lab^3^. The test scores loading reliably onto the verbal memory principal component included the Warrington Recognition Memory Test for Words^28^ and total learning and percent retained over 20 minutes from the Rey Auditory Verbal Learning Test^29^. The test scores that loaded reliably onto the visuospatial memory component included the Warrington Recognition Memory Test for Faces^28^, total learning on the Rey Visual Design Learning Test^30^, and trials to criterion performance on a spatial conditional associative learning task^31^. We transformed the raw pre- and post-surgical neuropsychological test scores of patients in the present study into summary verbal and visuospatial factor scores using the previously estimated factor loadings following previous work from our group^12,17,23,32^, in order to capture a more reliable representation of core abilities than afforded by single test scores. Preoperative verbal and visuospatial memory scores were then subtracted from postoperative scores to obtain measures of verbal and visuospatial memory change after surgery, which constitute the dependent variables in our models.

### MRI data acquisition

All structural and resting-state functional MRI scans were acquired preoperatively. For each subject, a high-resolution T1-weighted 3D anatomical scan (FOV 220 mm, 146 slices, 256×256 matrix, voxel size of 0.86 x 0.86 x 1.0) was collected on a 3T Signa MR system (GE Medical Systems, Milwaukee, WI). T2*-weighted resting-state fMRI scans were acquired with an echo-planar pulse imaging sequence (FOV 240 mm, 28-32 slices depending on head size, TR=2000 ms, TE=25 ms, 64×64 matrix, 3.75×3.75×5 mm voxels, 180 volumes). During the resting state scans, participants were instructed to lie still and “not to think about anything in particular” with their eyes closed.

### fMRI preprocessing

Preprocessing was performed in SPM8 (http://www.fil.ion.ucl.ac.uk/spm/software/spm8). The preprocessing pipeline included re-alignment, co-registration, normalization, temporal filtering for physiologic noise, and scrubbing of volumes with excess motion, as described in our prior publication^23^.

### Identification of the memory network and to-be-resected regions

We constructed a network of memory-relevant regions for resting-state analyses by selecting all of the regions from the Brainnetome Atlas (https://atlas.brainnetome.org/)^33^ that tended to be activated by tasks targeting explicit memory, episodic recall, or encoding, as identified using the meta data labels of the meta-analytical BrainMap Database associated with the atlas (http://www.brainmap.org/taxonomy). Contralateral homologues of the identified regions of interest (ROIs) were also included. Using this approach (which follows our previous work^23^), we obtained a total of 32 ROIs in each hemisphere (64 total) spanning frontal, temporal, and parietal regions, which we will refer to as the ‘memory network’ (visible in **Figure 2C**). We subsequently identified seven regions within the defined memory network in each temporal lobe that were within the limits of standard ATL resection^27^: medial area 38 (A38m) of the anterior superior temporal gyrus, rostral area 21 (A21r) of the anterior middle temporal gyrus, rostral area 35/36 of the parahippocampal gyrus (A35/36r), caudal area 35/36 of the parahippocampal gyrus (A35/36c), area 28/34 of the entorhinal cortex (A28/34), rostral hippocampus (rhipp), and caudal hippocampus (chipp). These 7 regions in the left hemisphere together comprise the ‘to-be-resected ATL’ in the left TLE group, while the homologous 7 regions in the right hemisphere comprise ‘the to-be-resected ATL’ in the right TLE group.

**Figure 1.**
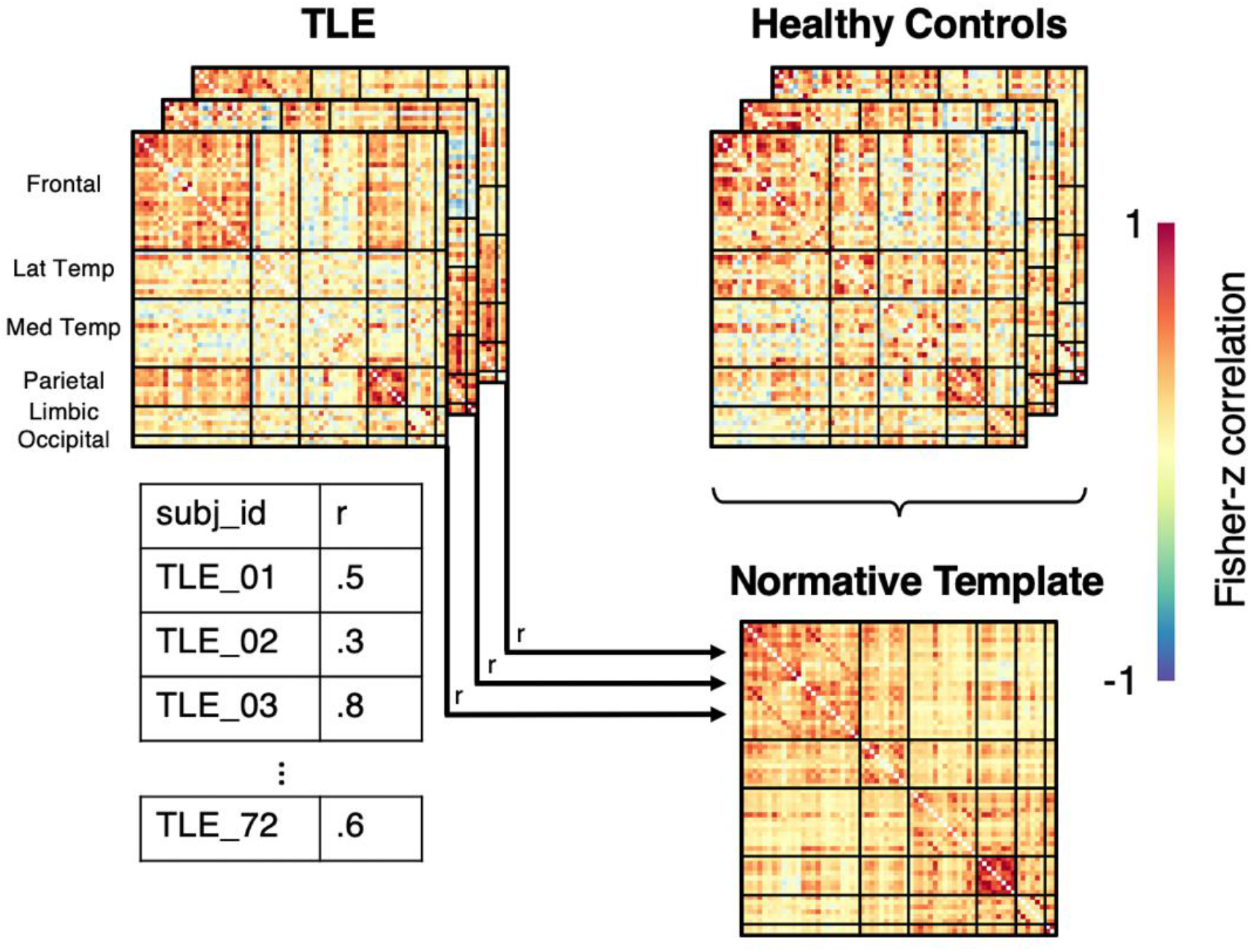
Matrix Similarity Derivation. Matrices representing each participant’s resting memory network connectivity were constructed. A normative template was created by averaging the matrices from 19 healthy controls, representative of typical memory network connectivity in the healthy brain. Each patient’s connectivity matrix was vectorized (not depicted here) and correlated with the (vectorized) normative template, producing a Pearson’s correlation coefficient representing matrix similarity for each patient. The resulting Pearson’s correlation coefficients were transformed using a Fisher’s z-transformation.

**Figure 2.**
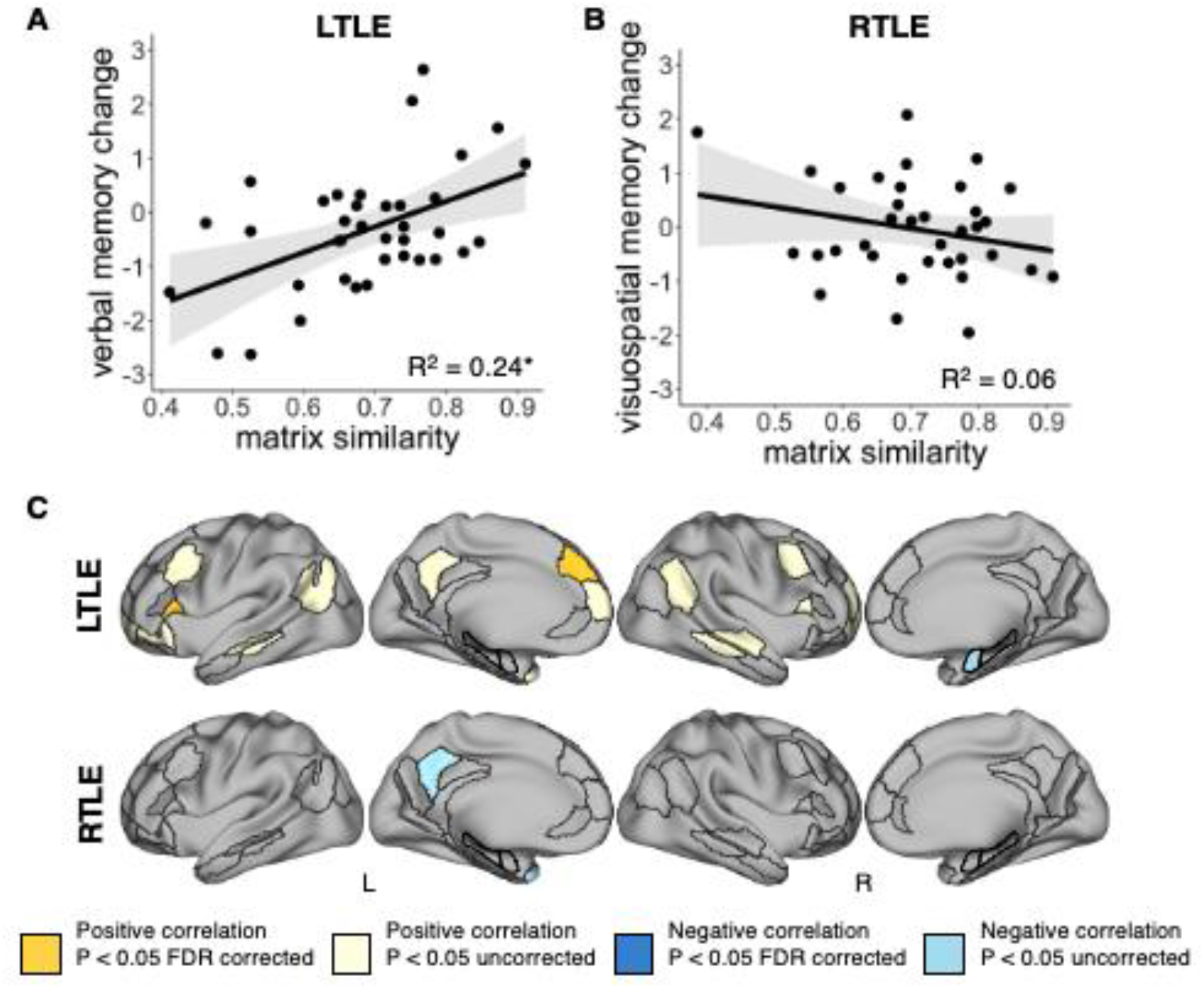
Matrix similarity and ROI-level similarity association with postoperative memory change. A) Matrix similarity correlation with postoperative verbal memory change in LTLE. B) Matrix similarity correlation with postoperative visuospatial memory change in RTLE. C) ROI-level similarity analysis results in patients with LTLE (top) and RTLE (bottom). Yellow areas denote regions where atypical preoperative connectivity was correlated with greater material-specific memory decline after surgery (positive correlations), and blue areas denote regions where normative preoperative connectivity was associated with greater material-specific memory decline after surgery (negative correlations). Darker colors represent ROIs that survive correction for multiple comparisons. L= left hemisphere, R=right hemisphere, FDR=False Discovery Rate, * denotes statistical significance p<0.05. Grey ribbons on correlation plots represent 95% confidence interval for a 2-tailed test.

### Matrix similarity analysis

We extracted the mean time course across all the unsmoothed voxels in each ROI of the memory network, which we correlated with the mean time course of every other ROI in a pairwise fashion. This resulted in a 64×64 connectivity matrix for each participant, representing subject-level memory network connectivity. We normalized the resulting Pearson’s correlation coefficients using a Fisher’s z-transformation. The control participants’ connectivity matrices were averaged together to create a normative template, representative of typical memory network connectivity in the healthy brain. We then correlated the vectorized normative healthy control matrix with each of the vectorized patient matrices, producing a Pearson’s correlation coefficient representing matrix similarity for each patient (**Figure 1**), which we normalized using a Fisher’s z-transformation. Higher matrix similarity values reflect a more normative pattern of connectivity. We then used linear regression to predict post-operative material-specific memory change (verbal memory for left TLE and visual memory for right TLE) using matrix similarity and group (left TLE/right TLE) as predictors.

### ROI similarity analysis

We additionally calculated ROI-level similarity between each patient’s network and the normative template to identify which ROIs’ pattern of connectivity were driving the overall relationship observed between matrix similarity and memory change. Specifically, we investigated each ROI’s connectivity fingerprint: it’s pattern of connectivity to every other ROI in the memory network^23^. We correlated each ROI’s connectivity fingerprint with that of the corresponding ROI of the normative template for each participant (i.e. correlating only corresponding rows in the control and patient matrices). This resulted in an ROI-level similarity value for each ROI in every patient, which represented how “normal” the pattern of connectivity of a given ROI was to the rest of the memory network. Higher ROI-level similarity values therefore reflect a more normative pattern of connectivity of a given ROI to every other ROI in the memory network (across both positive and negative connections). We Fisher z-transformed each correlation and evaluated the relationship between each region’s ROI-level similarity and postoperative memory change using Pearson’s correlations. We report uncorrected p-values <0.05 and p-values corrected for False Discovery Rate (FDR) across the 64 ROIs.

### Mean ATL degree analysis

We chose degree centrality – the number of connections a given region has to the rest of the network – as a relatively straightforward measure of regional functional integration, following our previous work^23^. Using the Brain Connectivity Toolbox (https://sites.google.com/site/bctnet/)^34^ and in-house scripts, individual subject memory network matrices were thresholded and binarized so that only the top positive connections survived at 5, 10, 15, 20, 25, 30, 35, 40, and 45 percentile connection density thresholds. By calculating several proportional thresholds, in contrast to absolute thresholds, we ensure that all participants’ matrices have similar number of edges, and reduce bias in selecting any one arbitrary threshold^34^. We next extracted degree centrality for each of the 7 to-be-resected ROIs at each connection density threshold for each patient. We summed the degree across the to-be-resected ROIs within each hemisphere and threshold, subtracting out the connections that link to-be-resected ROIs to each other. These summed degree measures served as our index of to-be-resected ATL integration with the rest of the memory network at each connection density threshold.

As statistical tests performed on neighbouring density thresholds are strongly dependent, corrections for multiple comparisons are not typically performed^35^. We therefore performed statistical tests on mean degree-centrality across the range of thresholds. To calculate mean degree, we first z-scored degree centrality within each threshold across groups to standardize degree scores. We then averaged the standardized scores across the range of thresholds for each subject. One-way ANOVAs were then conducted to test for differences in mean degree between groups (controls/left TLE/right TLE), and linear regression was used to predict material-specific postoperative memory change as a function of mean degree and group (left TLE/right TLE). The 8 participants who underwent selective amygdalohippocampectomy were excluded from this analysis, as the degree centrality of the to-be-resected regions was not directly comparable to those patients who underwent standard anterior temporal lobectomy.

### Combined Model Prediction

We performed multiple linear regression using the network measures that were reliably associated with material-specific memory change after surgery as predictors, to assess the unique influence of each variable on memory change. We ran a second linear regression that additionally included demographic and clinical variables of sex, education, age at preoperative assessment, age of onset, presence or absence of MTS, and preoperative material-specific memory capacity in the model. This allows a comparative assessment of unique and shared variance explained with network measures relative to conventional predictors of memory outcome.

### Data Availability

Deidentified memory network connectivity matrices for all participants, code used to calculate matrix similarity and mean ATL degree, and code for all statistical analyses and figures are available at https://github.com/saudrain/paper-MSMem2023.

## Results

### Participant demographics

Patient demographics can be observed in **Table 1**. We used one-way ANOVAs, t-tests, and chisquare tests to compare demographic variables between groups, and non-parametric Kruskal-Wallis and Wilcoxon rank sum tests where normality assumptions were violated. We found no significant differences in age between the three groups (χ^2^(2)=2.85, p=0.24; left TLE/right TLE/control), nor were there differences in sex (χ^2^(2)=2.57, p=0.28). However, there were differences in education (F(2,86)=12.83, p=0.000013) due to higher levels of education in controls than both patient groups (control vs left TLE: W=524.5, p=0.000082, MedianDiff=3, CI[2-5]; control vs right TLE: t(50)=4.33, p=0.000073, CI[1.95-5.33]). Years of education between the left and right TLE groups was not reliably different W=669.5, p=0.81, MedianDiff=0.00002, CI[-1-1]), nor was age of onset (W=558, p=0.32, MedianDiff=-3, CI[-11-3]). There were no reliable differences in hemisphere dominance (p=0.48, odds ratio=0.52, CI[0.077-2.71]), surgery type (p=0.47, odds ratio=1.87, CI[0.33-13.08]), or presence of MTS (χ^2^(1)=0.17, p=0.68) between the patient groups, nor were there differences in standardized preoperative material-specific memory scores (t(65.01)=-1.17, p=0.25, MeanDiff=0.34, CI[-0.90-0.23]) or material-specific memory change after surgery (t(68.8)=-1.22, p=0.23, MeanDiff=0.29, CI[-0.77-0.19]).

### Matrix Similarity of the memory network

A two sample t-test indicated that the distribution of matrix similarity values across left and right TLE groups was comparable (t(70)=-0.69, p=0.50, M left TLE=0.69, M right TLE=0.71, MeanDiff=-0.018, CI[-0.071-0.035]). We ran a linear regression predicting postoperative material-specific memory change (standardized verbal and visuospatial memory change scores for patients with left and right TLE respectively) using matrix similarity and group (left/right TLE) as predictors. We found significant effects of matrix similarity (β=4.73, CI[2-7.46], t(68)=3.46, p=0.0009), group (β=4.95, CI[2.12-7.79], t(68)=3.48, p=0.0009), and a significant interaction between the two (β=-6.73, CI[-10.75-(−2.7), t(68)=-3.33, p=0.001). A test of simple slopes found that the interaction was driven by a significant positive relationship between matrix similarity and material-specific memory change in the left TLE group (β=4.73, CI[2.01-7.46]) but not the right TLE group (β=-1.99, CI[-4.95-0.97]. Thus, left TLE patients who showed a pattern of preoperative memory network connectivity that was more dissimilar to that of the healthy control template showed the greatest decline in memory after surgery (**Figure 2.A**), but atypical connectivity was not reliably beneficial nor detrimental to memory after surgery in the right TLE group (**Figure 2.B**).

We examined ROI-level similarity to identify those regions contributing to the global effect seen in the left TLE group. Patients exhibiting more atypical preoperative patterns of connectivity in the left inferior frontal gyrus (caudal area 45/A45c; r=0.51, CI[0.23-0.72], p=0.04 FDR corrected) and left medial superior frontal gyrus (medial area 9/A9m; r=0.51, CI[0.22-0.72], p=0.04 FDR corrected) had worse verbal memory decline after surgery. Several other regions exhibited the same positive relationship with verbal memory change, but did not survive FDR correction (r>0.33; **Figure 2.C**). Nonetheless, the connectivity pattern of these regions likely contribute to the overall relationship between matrix similarity and verbal memory change after surgery, and suggests a bilateral contribution throughout much of the memory network. Only one ROI had a negative relationship between ROI similarity and verbal memory: the right anterior hippocampus, although this relationship did not survive corrections for multiple comparisons.

We additionally calculated an exploratory analysis of ROI-level similarity and its relationship with postoperative visuospatial memory change in right TLE, as we reasoned that the normative pattern of connectivity of fewer ROIs may associate with postoperative visuospatial memory change, and be masked by including the whole memory network in the analysis. We found a negative correlation in the left precuneus (area 31/A31; r=-0.36, CI[-0.62-(−0.02)], p=0.036) and left superior anterior temporal pole (medial area 38/A38m; r=−0.44, CI[−0.68-(−0.13)], p=0.007; **Figure 2.C**). In these regions, a normative pattern of connectivity was associated with worse visuospatial memory decline after surgery (the opposite pattern of that observed for left TLE patients). However, these correlations did not survive FDR correction.

### Degree centrality of to-be-resected regions

We next characterized group differences in functional integration of the to-be-resected region with the rest of the memory network. As the assumption of heterogeneity of variance was violated, we used a one-way Welch ANOVA and found a significant effect of group on mean degree centrality in the left ATL region (F(2,54.45)=8.16, p=0.0008). Post-hoc pairwise Grames Howell tests indicated that the left TLE group had lower mean degree centrality than right TLE (MeanDiff=0.55, CI[0.04-1.06], p=0.03) and control groups (MeanDiff=-0.87, CI[-1.38-(−0.35)], p=0.0005); **Figure 3.B**, left panel), but there was no significant difference between the right TLE and control groups (MeanDiff=-0.32, CI[-0.77-0.13], p=0.21). A similar analysis for the right ATL region demonstrated a significant effect of group (F(2,88)=11.69, p=0.000003), with lower mean degree centrality in right TLE compared to left TLE (MeanDiff=0.9, CI[0.51-1.3], t(88)=4.57, p<0.0001) and control groups (MeanDiff=0.81, CI[0.33-1.3], t(88)=3.37, p=0.001; **Figure 3.B,** right panel), and no difference between the left TLE and control groups (MeanDiff=-0.1, CI[-0.57-0.37], t(88)=0.41, p=0.68). Thus, both TLE groups demonstrated fewer positive functional connections between the epileptogenic ATL and the rest of the memory network, compared to non-epileptogenic or healthy ATLs.

**Figure 3.**
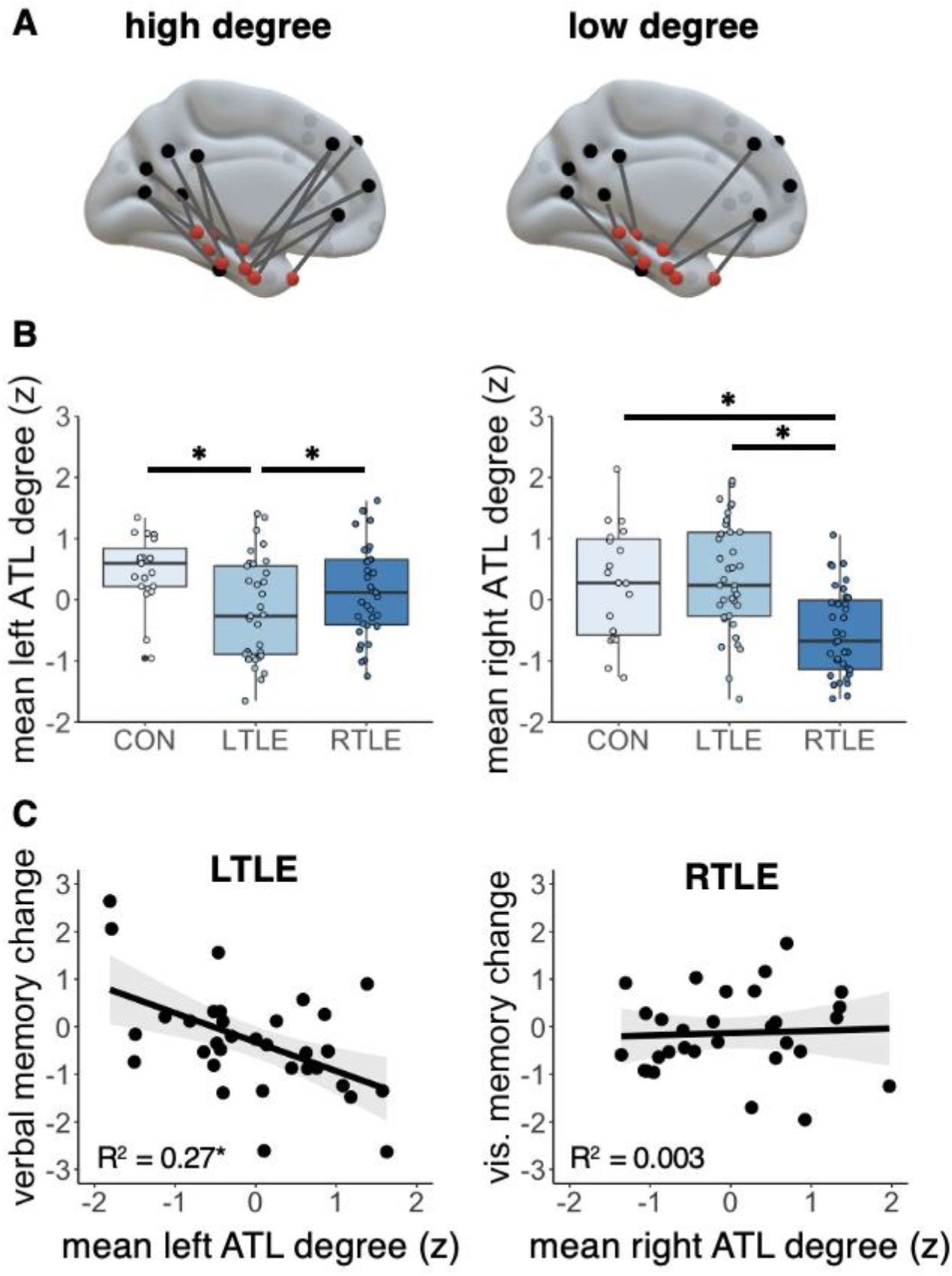
Mean ATL Degree and postoperative memory change. A) A schematic example of a brain with high and low degree centrality between the to-be-resected ATL (red nodes) and the rest of the memory network (black nodes). B) Group comparison of mean ATL degree between groups in the left hemisphere (to-be-resected region in the LTLE group; left plot), and the right hemisphere (to-be-resected region in the RTLE group; right plot). C) Correlation between mean ATL degree in the left hemisphere of the LTLE group and verbal memory change (left panel), and mean ATL degree in the right hemisphere of the RTLE group and visuospatial memory change (right panel). * denotes statistical significance p<0.05. Grey ribbons on correlation plots represent 95% confidence interval for a 2-tailed test.

To evaluate the contribution of this measure of functional integration to informing memory change, we conducted a linear model predicting standardized material-specific memory change (verbal memory in left TLE, visuospatial memory in right TLE) with mean ATL degree and group (left/right TLE) as predictors. We found a main effect of mean ATL degree (β=-0.57, CI[-0.89-(−0.25)], t(60)=-3.59, p=0.0007). There was no effect of group (β=0.36, CI[-0.15-0.86], t(60)=0.40, p=0.17), but there was a group by mean ATL degree interaction (β=0.63, CI[0.07-1.19], t(60)=2.26, p=0.027). An investigation of simple slopes for each group indicated a significant negative relationship between mean ATL degree and material-specific memory change in the left TLE group (β=-0.57, CI[-0.89-(–0.25)], **Figure 3.C**, left panel), but no reliable relationship in the right TLE group (β=0.060, CI[-0.40-0.52], **Figure 3.C**, right panel). The relationships between material-specific memory and mean ATL degree were consistent across connection density thresholds when visually inspected. These findings indicate that removal of dominant ATLs that are highly integrated with the rest of the memory network is detrimental to verbal memory, whereas this same relationship with visuospatial memory change was not observed after nondominant resection in the right TLE group.

### Combined regression models

As both matrix similarity and mean ATL degree were reliable predictors of verbal memory change in left TLE, we ran a multiple regression model to determine the unique variance in verbal memory change explained by each measure. The combined model explained 43.66% of variance in verbal memory change in left TLE (F(2,31)=12.01, p=0.0001). Of the total variance explained, matrix similarity explained 16.81% of unique variance (β=3.85, CI[1.25-6.44], t(31)=3.02, p=0.005), mean ATL degree explained 23.04% of unique variance (β=-0.53, CI[-0.83-(−0.22), t(31)=3.55, p=0.001), and 3.81% of the variance was shared between the two measures. Thus, both normativity of the overall network and the integration of to-be-resected region are unique explanatory factors, with a strong collective power in predicting verbal memory change following left ATL surgery.

Finally, we sought to determine if these measures added predictive value above and beyond conventional demographic and clinical variables that have previously been associated with verbal memory change. We re-ran the same linear regression model, this time additionally including age at preoperative assessment, sex, education, preoperative verbal memory scores, presence/absence of MTS and age of onset in the model. The model significantly predicted verbal memory change (F(8,25)=4.29, p=0.002, multiple R^2^=0.58 adjusted R^2^=0.44). Of the predictors, matrix similarity (β=3.77, CI[1.03-6.52], t(25)=2.83, p=0.009) and mean ATL degree (β=-0.55, CI[-0.86-(−0.24)], t(25)=-3.65, p=0.001) remained significant predictors of verbal memory change, continuing to explain 13.69% and 22.09% unique variance in verbal memory change respectively. The only other predictor reliably associated with verbal memory change in our sample was age of onset (β=-0.029, CI[−0.06-(−0.001)], t(25)=−2.17, p=0.040, semi-partial R^2^=0.078), although age at preoperative assessment was marginally positively associated with verbal memory change as well (β=0.028, CI[-0.002-0.06], t(25)=1.94, p=0.064, semi-partial R^2^=0.063). The rest of the predictors were not significant (all p >0.41). An F-test directly comparing regression models with and without inclusion of the clinical and demographic variables indicated that the demographic and clinical variables did not add predictive value to verbal memory change beyond the model including only the network measures (F(6,25)=1.40, p=0.25). A separate linear regression indicated that none of the clinical or demographic variables significantly predicted visuospatial memory change in right TLE (all p>0.17).

## Discussion

Similar to our prior examination of post-operative changes in lexical access^23^, here we demonstrate that rs-fMRI metrics of functional network integrity provide robust indicators of episodic memory decline following dominant anterior temporal lobe excision. In both cases, global similarity to network organization in controls (matrix similarity) and focal integration of subsequently excised tissue with the rest of the network (mean ATL degree) explained a large amount of unique variance in postoperative cognitive change. In the present case, these measures explained more variance in verbal memory change in our sample than age, sex, education, age of onset, presence or absence of MTS, or preoperative verbal memory scores. Importantly, both network measures can be accessed with a quick, task-free, resting state scan, which can be easily appended to language mapping tasks that are commonly collected in surgical candidates for little additional time and cost.

Notably, the relationship between matrix similarity and verbal memory decline in patients with left TLE was driven by abnormal connectivity patterns of regions broadly outside of the to-be-resected zone, with coupling patterns of the left IFG and left medial superior frontal gyrus supplying the strongest association. Two other studies similarly implicate resting left IFG coupling in postoperative language and verbal memory outcomes in left TLE^23,36^, suggesting that healthy coupling of this region is particularly important for preserving these cognitive faculties after surgery on the dominant hemisphere. Nonetheless, the contribution of other ROIs suggests that matrix similarity likely captures functional reserve subserved by multiple regions and highlights the value of considering the memory network beyond the temporal lobes during presurgical planning. In contrast to the functional reserve implied by this metric, the finding that greater integration of to-be-resected regions with the broader network (mean ATL degree) indicates greater memory decline after their removal is consistent with the pattern observed for other measures of functional adequacy, such as pre-operative memory scores and task-related activation. Of importance, these measures that leverage information about a broad intrinsic network of regions implicated in episodic encoding and retrieval appear to out-perform other variables that are commonly used as correlates or predictors of memory performance, which accords with other findings from our group^12,16^.

The observation that atypical preoperative memory networks are associated with greater verbal memory decline suggests that atypical networks are less resilient to focal network insult with surgery on the dominant hemisphere. While there is some evidence that reorganization could benefit preoperative language ability in left TLE^37–39^, evidence of a benefit of functional reorganization to memory is mixed. Some studies have documented adaptive recruitment of the unaffected MTL or connectivity with the contralateral hemisphere^14,40–42^ but this is contradicted by null effects in other studies^6,8,43–45^ or even a detrimental effect of atypical connectivity to memory capacity^12^. Irrespective of whether reorganization is adaptive for cognition preoperatively, the present results suggest that atypical networks are suboptimal for supporting memory functioning after surgical removal of key nodes in the ATL. We venture that healthy networks may contain more redundant or degenerate connections that allow functional compensation in the face of impairment in a subset of regions^46,47^. From this perspective, there are more opportunities to rely on the integrity of spared regions within a more typically configured memory network. In contrast, deviation away from normal network connectivity with persistent epileptic activity could restrict the ability to capitalize on redundant connections.

In contrast to the clear findings in left TLE patients, these measures were not sensitive to visuospatial memory change in patients with right TLE. In fact, none of the measures that we examined, including demographic and clinical variables, were related to visuospatial memory change in our right TLE group. Both material-specific baseline memory and change measures were comparable between the groups, suggesting that our measures of verbal and visuospatial memory were comparably sensitive and specific. A limitation of network measures is that including ROIs that are uninformative can add noise to the metric, so it’s possible that the network of memory regions chosen was not optimal to capture visuospatial memory capacity as assessed by standard neuropsychological tests. Given our past work showing an association between hippocampal-posterior cingulate cortex connectivity and visuospatial memory outcome in right TLE patients^16,17^, it is possible that a more restrictive network could improve sensitivity. Additionally, hemispheric differences in hippocampal functional connectivity with memory networks have been described in the healthy brain^11^. Inherent hemispheric differences in hippocampal-cortical network connectivity could engender different patterns of network disruption according to the lateralization of the epileptic focus. Indeed, a growing body of literature indicates different patterns of network connectivity in left and right TLE^13,15,48–50^, and matrix similarity is not sensitive to precisely *how* networks are atypical. We previously found different patterns of hippocampal-cortical connectivity in left and right TLE to be associated with material-specific memory^17^, and thus more work is needed to clarify the relationship between structure, function, and cognition in each group.

To conclude, while conventional measures utilized to predict postoperative memory capacity in patients with TLE largely focus on the integrity of the to-be resected region, our results suggest that consideration of the broader memory network is needed to improve predictions of verbal memory change in patients with left TLE. Atypical preoperative resting-state memory networks are associated with greater verbal memory decline after surgery on the dominant hemisphere, as is greater integration of to-be-resected tissue with the rest of the memory network. Our findings add to the growing body of evidence that low-frequency fluctuations at rest indicate intrinsic functional networks and thus have potential to be valuable for clinical decision making regarding cognitive outcomes following anterior temporal lobe resection.

## Acknowledgements

This work was funded by Ontario Brain Institute (EpLink). The authors would additionally like to thank Irene Giannoylis and Dr. David Gold for their assistance with data collection and organization.

## References

1. Sherman EMS, Wiebe S, Fay-Mcclymont TB, et al. Neuropsychological outcomes after epilepsy surgery: Systematic review and pooled estimates. Epilepsia. 2011;52(5):857–869. doi:10.1111/j.1528-1167.2011.03022.x

2. Harvey DJ, Naugle RI, Magleby J, et al. Relationship between presurgical memory performance on the Wechsler Memory Scale-III and memory change following temporal resection for treatment of intractable epilepsy. Epilepsy Behav. 2008;13(2):372–375. doi:10.1016/j.yebeh.2008.04.024

3. St-Laurent M, McCormick C, Cohn M, Mišić B, Giannoylis I, McAndrews MP. Using multivariate data reduction to predict postsurgery memory decline in patients with mesial temporal lobe epilepsy. Epilepsy Behav. 2014;31:220–227. doi:10.1016/j.yebeh.2013.09.043

4. Witt J-A, Coras R, Schramm J, et al. Relevance of hippocampal integrity for memory outcome after surgical treatment of mesial temporal lobe epilepsy. J Neurol. 2015;262(10):2214–2224. doi:10.1007/s00415-015-7831-3

5. Stasenko A, Kaestner E, Reyes A, et al. Association Between Microstructural Asymmetry of Temporal Lobe White Matter and Memory Decline After Anterior Temporal Lobectomy. Neurology. 2022;98(11):e1151–e1162. doi:10.1212/WNL.0000000000200047

6. Bonelli SB, Powell RHW, Yogarajah M, et al. Imaging memory in temporal lobe epilepsy: Predicting the effects of temporal lobe resection. Brain. 2010;133(4):1186–1199. doi:10.1093/brain/awq006

7. Powell HWR, Richardson MP, Symms MR, et al. Preoperative fMRI predicts memory decline following anterior temporal lobe resection. J Neurol Neurosurg Psychiatry. 2007;79(6):686–693. doi:10.1136/jnnp.2007.115139

8. Rabin ML. Functional MRI predicts post-surgical memory following temporal lobectomy. Brain. 2004;127(10):2286–2298. doi:10.1093/brain/awh281

9. Kramer MA, Cash SS. Epilepsy as a Disorder of Cortical Network Organization. Neurosci. 2012;18(4):360–372. doi:10.1177/1073858411422754

10. Barnett A, Audrain S, McAndrews MP. Applications of Resting-State Functional MR Imaging to Epilepsy. Neuroimaging Clin N Am. 2017;27(4):697–708. doi:10.1016/j.nic.2017.06.002

11. Barnett AJ, Reilly W, Dimsdale-Zucker HR, Mizrak E, Reagh Z, Ranganath C. Intrinsic connectivity reveals functionally distinct cortico-hippocampal networks in the human brain. Kaplan R, ed. PLOS Biol. 2021;19(6):e3001275. doi:10.1371/journal.pbio.3001275

12. McCormick C, Protzner AB, Barnett AJ, Cohn M, Valiante TA, McAndrews MP. Linking DMN connectivity to episodic memory capacity: What can we learn from patients with medial temporal lobe damage? NeuroImage Clin. 2014;5:188–196. doi:10.1016/j.nicl.2014.05.008

13. Roger E, Pichat C, Torlay L, et al. Hubs disruption in mesial temporal lobe epilepsy. A resting-state fMRI study on a language-and-memory network. Hum Brain Mapp. 2020;41(3):779–796. doi:10.1002/hbm.24839

14. Holmes M, Folley BS, Sonmezturk HH, et al. Resting state functional connectivity of the hippocampus associated with neurocognitive function in left temporal lobe epilepsy. Hum Brain Mapp. 2014;35(3):735–744. doi:10.1002/hbm.22210

15. Doucet G, Osipowicz K, Sharan A, Sperling MR, Tracy JI. Extratemporal functional connectivity impairments at rest are related to memory performance in mesial temporal epilepsy. Hum Brain Mapp. 2013;34(9):2202–2216. doi:10.1002/hbm.22059

16. McCormick C, Quraan M, Cohn M, Valiante TA, McAndrews MP. Default mode network connectivity indicates episodic memory capacity in mesial temporal lobe epilepsy. Epilepsia. 2013;54(5):809–818. doi:10.1111/epi.12098

17. Barnett AJ, Man V, McAndrews MP. Parcellation of the hippocampus using resting functional connectivity in temporal lobe epilepsy. Front Neurol. 2019;10:1–12. doi:10.3389/fneur.2019.00920

18. Ives-Deliperi V, Butler JT. Mechanisms of cognitive impairment in temporal lobe epilepsy: A systematic review of resting-state functional connectivity studies. Epilepsy Behav. 2021;115:107686. doi:10.1016/j.yebeh.2020.107686

19. Doucet GE, He X, Sperling M, Sharan A, Tracy JI. Gray Matter Abnormalities in Temporal Lobe Epilepsy: Relationships with Resting-State Functional Connectivity and Episodic Memory Performance. Biagini G, ed. PLoS One. 2016;11(5):e0154660. doi:10.1371/journal.pone.0154660

20. Audrain S, McAndrews MP. Cognitive and functional correlates of accelerated long-term forgetting in temporal lobe epilepsy. Cortex. 2019;110:101–114. doi:10.1016/j.cortex.2018.03.022

21. Fleury M, Buck S, Binding LP, et al. Episodic memory network connectivity in temporal lobe epilepsy. Epilepsia. 2022;63(10):2597–2622. doi:10.1111/epi.17370

22. McAndrews MP, Barnett A. Clinical Utility of Resting State Functional MRI. In: Habas C, ed. The Neuroimaging of Brain Diseases: Structural and Functional Advances. Springer International Publishing; 2018:59–79. doi:10.1007/978-3-319-78926-2_3

23. Audrain S, Barnett AJ, McAndrews MP. Language network measures at rest indicate individual differences in naming decline after anterior temporal lobe resection. Hum Brain Mapp. 2018;39:4404–4419. doi:10.1002/hbm.24281

24. Dulay MF, Busch RM. Prediction of neuropsychological outcome after resection of temporal and extratemporal seizure foci. Neurosurg Focus. 2012;32(3):E4. doi:10.3171/2012.1.FOCUS11340

25. Binder JR, Sabsevitz DS, Swanson SJ, Hammeke TA, Raghavan M, Mueller WM. Use of preoperative functional MRI to predict verbal memory decline after temporal lobe epilepsy surgery. Epilepsia. 2008;49(8):1377–1394. doi:10.1111/j.1528-1167.2008.01625.x

26. Busch RM, Hogue O, Miller M, et al. Nomograms to Predict Verbal Memory Decline After Temporal Lobe Resection in Adults With Epilepsy. Neurology. 2021;97(3):e263–e274. doi:10.1212/WNL.0000000000012221

27. Mansouri A, Fallah A, McAndrews MP, et al. Neurocognitive and Seizure Outcomes of Selective Amygdalohippocampectomy versus Anterior Temporal Lobectomy for Mesial Temporal Lobe Epilepsy. Epilepsy Res Treat. 2014;2014:1–8. doi:10.1155/2014/306382

28. Warrington E. Recognition Memory Test: Manual. NFER-Nelson; 1984.

29. Strauss E, Sherman EMS, Spreen OA. Compendium of Neuropsychological Tests: Administration, Norms, and Commentary. 3rd ed. Oxford University Press; 2006.

30. Spreen O, Strauss E. A Compendium OfNeuropsychological Tests: Administration, Norms and Commentary. Oxford University Press; 1991.

31. Petrides M. Deficits on conditional associative-learning tasks after frontal- and temporal lobe lesions in man. Neuropsychologia. 1985;23(5):601–614. doi:10.1016/0028-3932(85)90062-4

32. Barnett AJ, Park MTM, Pipitone J, Chakravarty MM, McAndrews MP. Functional and structural correlates of memory in patients with mesial temporal lobe epilepsy. Front Neurol. 2015;6(MAY):1–9. doi:10.3389/fneur.2015.00103

33. Fan L, Li H, Zhuo J, et al. The Human Brainnetome Atlas: A New Brain Atlas Based on Connectional Architecture. Cereb Cortex. 2016;26(8):3508–3526. doi:10.1093/cercor/bhw157

34. Rubinov M, Sporns O. Complex network measures of brain connectivity: Uses and interpretations. Neuroimage. 2010;52(3):1059–1069. doi:10.1016/j.neuroimage.2009.10.003

35. Fornito A, Zalesky A, Breakspear M. Graph analysis of the human connectome: Promise, progress, and pitfalls. Neuroimage. 2013;80:426–444. doi:10.1016/j.neuroimage.2013.04.087

36. Doucet G, Rider R, Taylor N, et al. Presurgery resting-state local graph-theory measures predict neurocognitive outcomes after brain surgery in temporal lobe epilepsy. Epilepsia. 2015;56(4):517–526. doi:10.1111/epi.12936

37. Berl MM, Balsamo LM, Xu B, et al. Seizure focus affects regional language networks assessed by fMRI. Neurology. 2005;65(10):1604–1611. doi:10.1212/01.wnl.0000184502.06647.28

38. Brázdil M, Zákopčan J, Kuba R, Fanfrdlová Z, Rektor I. Atypical hemispheric language dominance in left temporal lobe epilepsy as a result of the reorganization of language functions. Epilepsy Behav. 2003;4(4):414–419. doi:10.1016/S1525-5050(03)00119-7

39. Weber B, Wellmer J, Reuber M, et al. Left hippocampal pathology is associated with atypical language lateralization in patients with focal epilepsy. Brain. 2006;129(2):346–351. doi:10.1093/brain/awh694

40. Figueiredo P, Santana I, Teixeira J, et al. Adaptive visual memory reorganization in right medial temporal lobe epilepsy. Epilepsia. 2008;49(8):1395–1408. doi:10.1111/j.1528-1167.2008.01629.x

41. Li H, Ji C, Zhu L, et al. Reorganization of anterior and posterior hippocampal networks associated with memory performance in mesial temporal lobe epilepsy. Clin Neurophysiol. 2017;128(5):830–838. doi:10.1016/j.clinph.2017.02.018

42. Cheung M-C, Chan AS, Lam JMK, Chan Y-L. Pre- and postoperative fMRI and clinical memory performance in temporal lobe epilepsy. J Neurol Neurosurg Psychiatry. 2009;80(10):1099–1106. doi:10.1136/jnnp.2009.173161

43. Powell HWR, Richardson MP, Symms MR, et al. Reorganization of Verbal and Nonverbal Memory in Temporal Lobe Epilepsy Due to Unilateral Hippocampal Sclerosis. Epilepsia. 2007;48(8):1512–1525. doi:10.1111/j.1528-1167.2007.01053.x

44. Bonelli SB, Thompson PJ, Yogarajah M, et al. Memory reorganization following anterior temporal lobe resection: a longitudinal functional MRI study. Brain. 2013;136(6):1889–1900. doi:10.1093/brain/awt105

45. Limotai C, McLachlan RS, Hayman-Abello S, et al. Memory loss and memory reorganization patterns in temporal lobe epilepsy patients undergoing anterior temporal lobe resection, as demonstrated by pre-versus post-operative functional MRI. J Clin Neurosci. 2018;55:38–44. doi:10.1016/j.jocn.2018.06.020

46. Tononi G, Sporns O, Edelman GM. Measures of degeneracy and redundancy in biological networks. ProcNatlAcadSci,USA. 1999;96(6):3257–3262. doi:10.1073/pnas.96.6.3257

47. Aerts H, Fias W, Caeyenberghs K, Marinazzo D. Brain networks under attack: robustness properties and the impact of lesions. Brain. 2016;139(12):3063–3083. doi:10.1093/brain/aww194

48. Besson P, Dinkelacker V, Valabregue R, et al. Structural connectivity differences in left and right temporal lobe epilepsy. Neuroimage. 2014;100:135–144. doi:10.1016/j.neuroimage.2014.04.071

49. Pittau F, Grova C, Moeller F, Dubeau F, Gotman J. Patterns of altered functional connectivity in mesial temporal lobe epilepsy. Epilepsia. 2012;53(6):1013–1023. doi:10.1111/j.1528-1167.2012.03464.x

50. Chiang S, Stern JM, Engel J, Levin HS, Haneef Z. Differences in graph theory functional connectivity in left and right temporal lobe epilepsy. Epilepsy Res. 2014;108(10):1770–1781. doi:10.1016/j.eplepsyres.2014.09.023

